# Brain age as an estimator of neurodevelopmental outcome: A deep learning approach for neonatal cot-side monitoring

**DOI:** 10.1101/2023.01.24.525361

**Authors:** Amir Ansari, Kirubin Pillay, Luke Baxter, Emad Arasteh, Anneleen Dereymaeker, Gabriela Schmidt Mellado, Katrien Jansen, Gunnar Naulaers, Aomesh Bhatt, Sabine Van Huffel, Caroline Hartley, Maarten De Vos, Rebeccah Slater

**Affiliations:** Department of Electrical Engineering (ESAT), STADIUS Center for Dynamical Systems, Signal Processing and Data Analytics, KU Leuven, Leuven, Belgium; Department of Paediatrics, University of Oxford, Oxford, UK; Department of Neonatology, Wilhelmina Children’s Hospital, University Medical Center Utrecht, Utrecht, the Netherlands; Department of Development and Regeneration, University Hospitals Leuven, Neonatal Intensive Care Unit, KU Leuven, Leuven, Belgium; Department of Development and Regeneration, University Hospitals Leuven, Child Neurology, KU Leuven, Leuven, Belgium

**Keywords:** Preterm, Electroencephalography, Machine Learning, Artificial Intelligence, Convolutional Neural Network, Bayley Scales

## Abstract

The preterm neonate can experience stressors that affect the rate of brain maturation and lead to long-term neurodevelopmental deficits. However, some neonates who are born early follow normal developmental trajectories. Extraction of data from electroencephalography (EEG) signals can be used to calculate the neonate’s brain age which can be compared to their true age. Discrepancies between true age and brain age (the brain age delta) can then be used to quantify maturational deviation, which has been shown to correlate with long-term abnormal neurodevelopmental outcomes. Nevertheless, current brain age models that are based on traditional analytical techniques are less suited to clinical cot-side monitoring due to their dependency on long-duration EEG recordings, the need to record activity across multiple EEG channels, and the manual calculation of predefined EEG features which is time-consuming and may not fully capture the wealth of information in the EEG signal. In this study, we propose an alternative deep-learning approach to determine brain age, which operates directly on the EEG, using a Convolutional Neural Network (CNN) block based on the Inception architecture (called Sinc). Using this deep-learning approach on a dataset of preterm infants with normal neurodevelopmental outcomes (where we assume brain age = postmenstrual age), we can calculate infant brain age with a Mean Absolute Error (MAE) of 0.78 weeks (equivalent to a brain age estimation error for the infant within +/− 5.5 days of their true age). Importantly, this level of accuracy can be achieved by recording only 20 minutes of EEG activity from a single channel. This compares favourably to the degree of accuracy that can be achieved using traditional methods that require long duration recordings (typically >2 hours of EEG activity) recorded from a higher density 8-electrode montage (MAE = 0.73 weeks). Importantly, the deep learning model’s brain age deltas also distinguish between neonates with normal and severely abnormal outcomes (Normal MAE = 0.71 weeks, severely abnormal MAE = 1.27 weeks, p=0.02, one-way ANOVA), making it highly suited for potential clinical applications. Lastly, in an independent dataset collected at an independent site, we demonstrate the model’s generalisability in age prediction, as accurate age predictions were also observed (MAE of 0.97 weeks).

**Highlights:**

- Preterm stress exposure leads to long-term neurodevelopmental deficits
- Deficits are quantifiable using EEG-based brain age prediction errors
- Our deep-learning solution for brain age prediction outperforms previous approaches
- Predictions are achieved with only 20 mins EEG and a single bipolar channel
- Prediction errors correlate with long-term Bayley scale neurodevelopmental outcomes

## 1. Introduction

The newborn infant’s brain is undergoing rapid developmental change, influenced by both genetic and environmental factors (Colonnese et al., 2010; Milh et al., 2007; Wess et al., 2017). Relative to their term-born counterparts, infants born prematurely are at increased risk of poorer long-term neurodevelopmental outcomes (Blencowe et al., 2013; Wallois et al., 2020). This risk of impairment increases with the degree of prematurity at birth and the presence of gross morphological lesions, but can also be brought about by subtler environmental stressors (Scher, 2008), excessive exposure to painful stimuli (Grunau, 2013; Moultrie et al., 2017), and pharmacological interventions (Duerden et al., 2016; Malk et al., 2014).

The early identification of abnormal neurodevelopment is essential to identify infants at greatest risk who might benefit most from developmental care interventions (Burke, 2018). To date, neurological assessment of the newborn has remained predominantly subjective (Dempsey et al., 2018). For example, trained neonatologists and clinical neurophysiologists visually inspect infant’s brain activity using electroencephalography (EEG) to determine if brain function is developmentally age-appropriate or dysmature (Scher, 1997), based on developmentally changing EEG features characteristic of maturational status (André et al., 2010). While these trained individuals can estimate age with an error of two weeks for preterm babies and one week for term babies, these estimates can be highly variable across reviewers (Stevenson et al., 2020b). Subjectivity, inter-rater variability, and requirement of specialist EEG interpretation are central issues that severely limit the reliability and generalisability of many current neurological assessment methods. There is an urgent need for objective and automated neuromonitoring that can be used cot-side to identify infants at increased risk of abnormal neurodevelopmental outcomes.

To this end, a variety of metrics have been developed to capture key maturational characteristics from the preterm EEG (De Wel et al., 2017; Dereymaeker et al., 2016; Lavanga et al., 2017; Pillay et al., 2018; Tolonen et al., 2007), and these measures have been combined using machine learning algorithms to successfully predict infants’ brain age (O’Toole et al., 2016; Stevenson et al., 2017). An infant’s brain age is their predicted age from a model that has been trained using brain-based features (structural or functional) as predictors and true age as the response. In adults, the difference between the brain age and the true age, termed the brain age delta, has been demonstrated to be more than random noise prediction error, but in fact is of biological and clinical value (Smith et al., 2019; Vidal-Pineiro et al., 2021).

In infants, analogous findings have been observed. Recently, we trained a Random Forest (RF) regression model using a data-driven approach that combined 226 EEG features and demonstrated a significant correlation between the infants’ brain age delta and the severity of their abnormal neurodevelopmental outcome, where the neurodevelopmental outcomes were assessed behaviourally using the Bayley Scales of Infant Development (BSID-II) at a 9-month follow-up test occasion (Pillay et al., 2020). Additionally, an independent research group showed a similar correlation when training a multivariate regression model for brain age estimation (Stevenson et al., 2020a). These studies established the proof-of-concept in infant populations that the inter-individual variability in automatically and objectively generated brain age deltas could be used to risk-stratify infants in the first few weeks of postnatal life according to neurodevelopmental outcomes.

However, a major limitation to these studies is their lack of clinical utility. A large number of features are required to summarize the EEG data, which are computationally time-consuming to calculate. These approaches rely on pre-staging the EEG recording into sleep states (i.e. sleep-staging) or burst periods which require additional algorithms (Dereymaeker et al., 2017b; Palmu et al., 2010). Furthermore, multiple EEG channels are required as well as at least 1 hour EEG recording duration. These data-heavy requirements severely limit the ease with which these methods can be incorporated into the busy clinical environment.

Here, we directly address these barriers to clinical utility by adopting a deep learning approach. Deep learning has demonstrated superior performance over traditional machine learning methods, has excellent performance on a reduced number of EEG channels, and tends to perform predictions faster once trained (Ansari et al., 2018). Furthermore, deep learning models are gaining popularity in preterm EEG analysis for classifying seizures (Ansari et al., 2019; O’Shea et al., 2021) and for automated sleep-staging (Ansari et al., 2020). Together, these observations suggest deep learning could offer a promising approach for cot-side monitoring and assessment of neurological function.

In the current study, we implement a novel Convolutional Neural Network (CNN)-based architecture, inspired by Google’s Inception model (and its variants), to generate infant brain age predictions using dramatically reduced EEG data requirements compared to previous proof-of-concept studies. We use our established RF model as a “gold standard” benchmark of performance, a model which requires eight EEG channels, at least 1 hour EEG recording duration, and EEG data sleep-staging. We train the RF and deep learning models on a training dataset, and subsequently test the models’ performance on two independent datasets, demonstrating robust external validation. Using our deep learning approach, we achieve performance comparable to our RF model benchmark, while requiring only a single EEG channel (1-channel bipolar montage), 20 mins EEG recording duration, and no EEG data sleep-staging. Our deep learning model is able to accurately predict infant age within the first few weeks of postnatal life, and generates brain age deltas with magnitudes that significantly differ between infants with normal and severely abnormal neurodevelopmental outcomes assessed using BSID-II at 9-month follow-up. This study thus demonstrates potential clinical utility for an objective and automated deep learning-based approach to cot-side assessment of infants’ neurological function and neurodevelopmental outcomes.

## 2. Methods

### 2.1. Participants

#### 2.1.1. Study Design

Data were collected in three independent cohorts. The first cohort, referred to as dataset 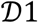, was used to train the models and compare the relative performances among models e.g. models with different architectures, different channel montages, and different recording durations. The second cohort, referred to as dataset 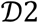, was used to independently test the trained RF and deep learning models in their brain age prediction performances, and to assess the association between brain age deltas and 9-month BSID-II follow-up outcomes. The third cohort, referred to as dataset 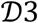, was used to further test the generalisability of the deep learning model to predict brain age in this dataset collected at an independent site by an independent research team.

#### 2.1.2. Recruitment

EEG data for datasets 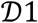 and 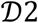 were recorded from the Neonatal Intensive Care Unit (NICU) at UZ Leuven Hospitals, Leuven, Belgium. Infants were recruited and data recorded with informed consent from the parents and in accordance with the guidelines approved by the ethics committee of the University Hospitals, Leuven. All infants had a gestational age (GA) at birth less than 32 weeks, and between two and four recordings were obtained during their stay in the NICU.

Infants in dataset 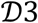 were selected from a database of previously recorded data collected at the Newborn Care Unit and Maternity wards of the John Radcliffe Hospital (Oxford University Hospitals NHS Foundation Trust, Oxford, United Kingdom). Ethical approval was obtained from the UK National Research Ethics Service (reference: 12/SC/0447) and parental written informed consent was obtained before each participant was studied.

All participant recruitment was conducted in accordance with the standards set by the Declaration of Helsinki and Good Clinical Practice guidelines.

#### 2.1.3. Datasets

Datasets 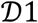 and 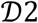 were collected as previously described (Pillay et al., 2020). Dataset 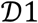 consists of n=40 infants (111 recordings) with postmenstrual age range (PMA) at time of recording of 27.3–43.1 weeks, with mean recording duration of 8h 07m (standard deviation: 5h 55m) and mean number of recordings per infant of 2.8 (standard deviation: 1.6). All infants in dataset 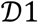 were selected for normal neurodevelopmental outcome at 24-months follow-up age based on behavioural assessment using BSID-II.

Dataset 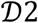 consists of n=43 infants (142 recordings). One infant with a single recording was excluded as our objective with this dataset was to assess longitudinal multi-recording trajectories. The analysed dataset 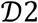 thus consists of n=42 infants (141 recordings) with a PMA at recording range of 27.3–42.0 weeks, mean recording duration of 7h 05m (standard deviation: 5h 43m), and mean number of recordings per infants of 3.3 (standard deviation: 1.4). Unlike dataset 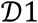, dataset 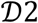 includes infants with a range of both normal and abnormal outcomes, grouped by BSID-II scores at 9-month follow-up (Pillay et al., 2020). N=22 infants (71 recordings) had normal outcome i.e. no neurodevelopmental impairment (NDI); n=10 infants (36 srevoereinNgDsI)) ohraddiedm(ilPdasacablneotrmal.a,l20o2u0tc).ome (mild NDI); and n=10 infants (34 recordings) had moderate-to-severe abnormal outcome (mild-to-severe NDI) or died (Pascal et al., 2020).

Dataset 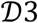 consists of n=73 infants, each recorded on a single test occasion (thus 73 recordings). Infants were included in this dataset for the current study if they had at least 20 minutes of EEG data recorded and if the EEG was assessed as normal for age by a trained clinical neurophysiologist (author GSM). The infants had a median PMA at recording of 35.3 weeks (interquartile range: 33.3 – 36.9, range: 28.0 – 42.6) and postnatal age of 14 days (interquartile range: 5 – 41, range: 0 – 112). The mean recording duration was 50 minutes (standard deviation: 18 minutes).

### 2.2. EEG data

#### 2.2.1. Setup

For dataset 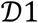 and 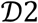, data were recorded using a sampling frequency of 250 Hz using Brain RT OSG Equipment (Mechelen, Belgium). In a few cases, the EEG was sampled at 256 Hz due to some setup variations on the Brain RT device used. All recordings were performed with nine electrodes in a referential montage: Fp1, Fp2, C3, C4, T3, T4, O1, O2, and Cz reference (Figure 1).

**Figure 1:**
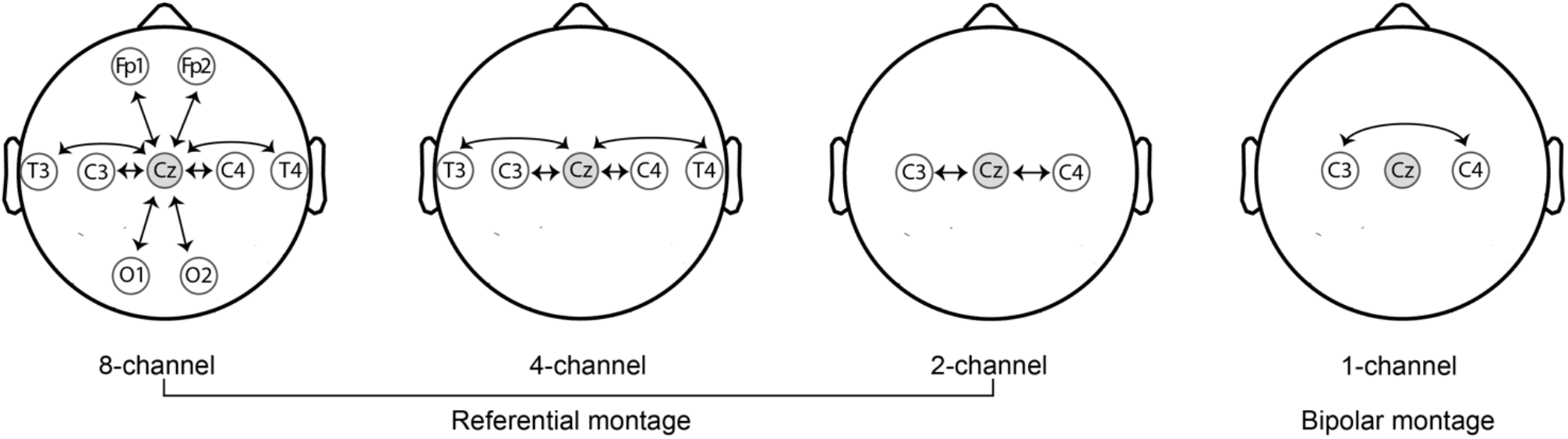
EEG montages used during analysis. All recordings in datasets 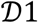 *and* 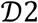 were acquired with eight recording EEG electrodes in positions: Fp1, Fp2, C3, C4, T3, T4, O1, O2, and a reference electrode placed at Cz (shaded in grey). The arrows represent the specific channels used during analysis. *For dataset* 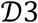, analysis was conducted using the 1-channel bipolar montage. Recordings were initially acquired with electrode positions Cz, CPz, C3, C4, Oz, FCz, T3 and T4, and a reference electrode at Fz.

For dataset 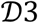, EEG recordings were acquired from DC to 800 Hz using a SynAmps RT 64-channel headbox and amplifiers (Compumedics Neuroscan). Activity was recorded using CURRY scan7 neuroimaging suite (Compumedics Neuroscan), with a sampling rate of 2000 Hz. Between 8 and 25 electrodes were used for recording, positioned according to the modified international 10-20 system, including C3 and C4 (those used in the analysis here), with reference at Fz and ground at Fpz. The scalp was cleaned with preparation gel (Nuprep gel, D.O. Weaver and Co.) and disposable Ag/AgCl cup electrodes (Ambu Neuroline) were placed with conductive paste (Elefix EEG paste, Nihon Kohden).

#### 2.2.2. Preprocessing

For the deep learning approaches in datasets 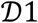 and 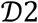, each recording was downsampled to 64 Hz to reduce the number of parameters required to train the model. The downsampling routine included pre-filtering to prevent aliasing using a low-pass filter with cut-off frequency 32 Hz. Filtering and downsampling was performed using the scipy.signal.resample_poly function. Recordings were then split into 30-second segments and the amplitudes standardized such that the mean and standard deviation of the amplitudes were zero and one, respectively. The mean and standard deviation were obtained by standardizing the data (across all channels) in the training set, with these values carried forward to standardize the test sets (see below). Finally, any segments where the absolute differences (compared to the mean) at any point exceeded 600 μV were rejected as artefact.

For the RF approach in datasets 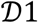 and 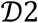, which relied on an explicit pre-calculation of many established features, the pre-processing approach (resampling and standardization) was different and specific to each calculated feature, as described previously (Pillay et al., 2018).

For dataset 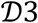, pre-processing was matched to the 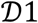 and 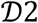 deep learning approach. We applied a low-pass 32 Hz anti-aliasing filter followed by downsampling to 64 Hz. For standardization of dataset 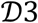, the mean and standard deviation of 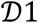 were used.

### 2.3. Brain age prediction architectures

#### 2.3.1. Sinc architecture

Figure 2a shows the block-diagram of the proposed deep neural network for brain age prediction. As input, the network processes a 30 s multi-channel EEG segment. Each input segment has dimension *C* × 1920 where *C* is the number of EEG channels and 1920 is the total number of timepoints in the 30 s segment (30 s duration × 64 Hz sampling frequency). Each segment has a single output label that is a continuous PMA value.

**Figure 2:**
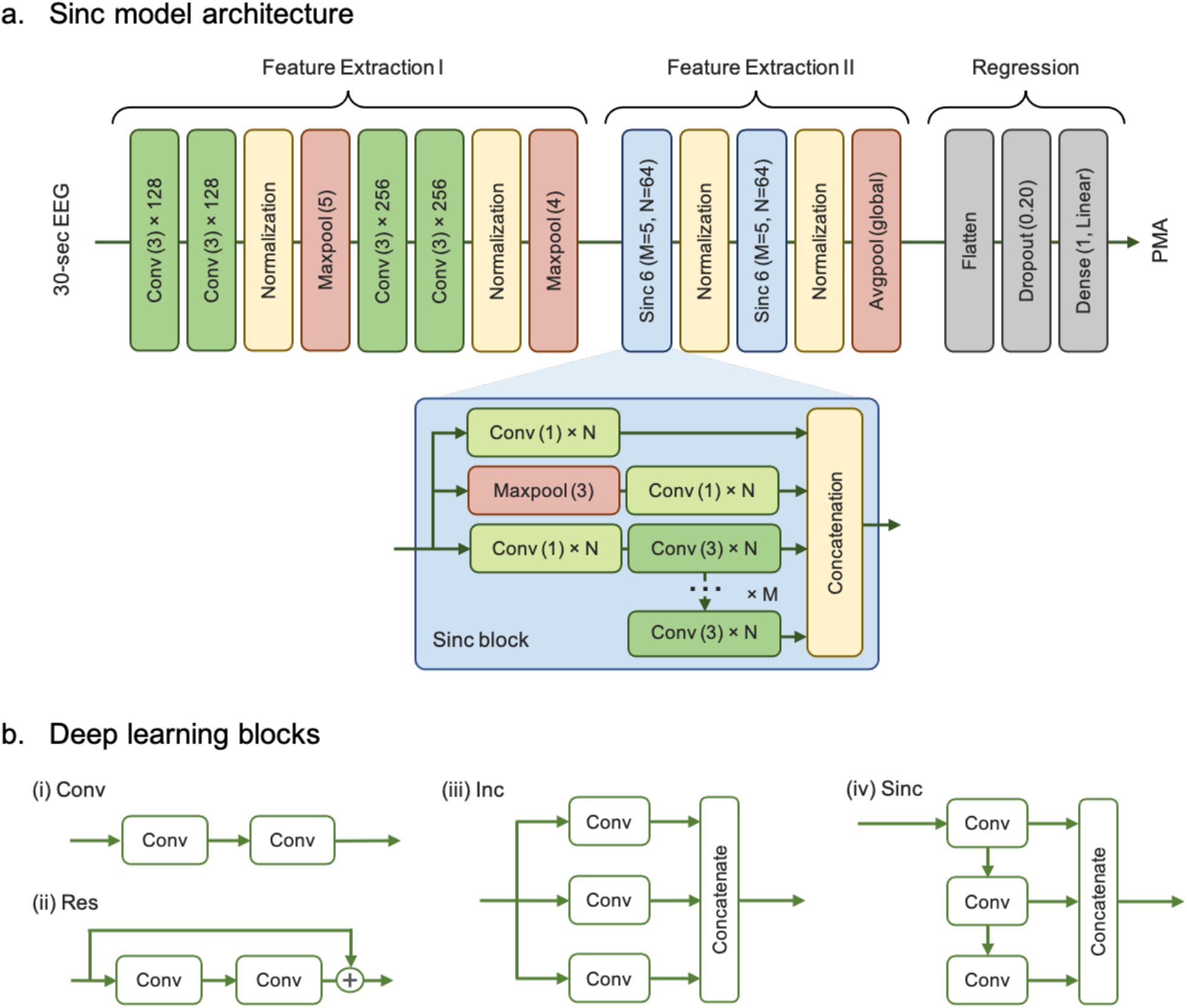
Deep learning architectures. **a.** Block-diagram of the proposed Sinc network architecture, including the typical structure of the Sinc block. **b.** Illustrative block diagrams of different blocks in the deep architectures: (i) Sequential Convolutional layers, (ii) Residual block, (iii) Inception block, (iv) Shared Inception (Sinc) block.

The model includes a series of convolutional layers with exponential linear unit (ELU) activations, maximum and average pooling layers to downsample the data, normalization layers for faster training convergence, and a dense layer with linear activation to perform the final regression and produce a brain age estimate. As each convolutional layer is designed to extract specific characteristics from the EEG, these are analogous to a (trainable, data-driven) feature extraction layer. More generally, the proposed architecture can be grouped into a more traditional, sequential CNN block that can be described as an initial feature extraction stage, followed by the two successive Sinc (i.e. Shared Inception) blocks that form a second feature extraction stage.

We previously introduced Sinc as a powerful CNN-based block for extracting multi-scale temporal information from infant EEG, namely sleep state classification (Ansari et al., 2021). Sinc is an extension of Google’s Inception block, where the original independent and parallel convolutional branches are now boosted via parameter sharing. As shown in Figure 2b, the output from each preceding branch is additionally fed into the subsequent one, with the overall output of Sinc comprising the concatenation of all multi-scale convolutions in the block (see also Figure 2biv). This increases the number of temporal scales achievable (by allowing a wider range of receptive fields), when compared to an Inception layer, while avoiding the need to scale up the number of trainable parameters as a result. Only two hyper-parameters are required for a Sinc block: *M* (the number of convolutional branches), and *N* (the number of convolutional filters used in each branch). When using a single-channel EEG segment as input (*C* = 1), the total number of trainable parameters in the complete model is 620K.

#### 2.3.2. Alternative deep learning architectures

Four different deep learning architectures were also considered, based on recent key developments in the CNN domain (Figure 2b), and the Sinc model was compared against these architectures: No FEII (following the same design as the Sinc architecture but without the entire Feature Extraction II portion), CNN (replaces the Sinc blocks with the same convolutional neural network layer used elsewhere in the model), Residual (similar to CNN but including the additional residual shortcut), and Inc (replacing the Sinc blocks with traditional Inception blocks). These architectures are described in Supplementary Information S.1.

#### 2.6.2. Random Forest (RF) architecture

In addition to comparing the Sinc architecture to alternative deep neural network architectures, it was also compared against our established and previously published RF approach (Pillay et al., 2020). The RF model was developed using a large set of pre-calculated features, derived after the EEG is classified into different sleep stages using an additional, unsupervised algorithm known as Cluster-based Adaptive Sleep Staging (CLASS) that we have also previously developed (Dereymaeker et al., 2017b; Pillay et al., 2020). The pre-calculated features were derived from an EEG literature review covering the amplitude domains, Fourier transforms, Wavelet transforms, Empirical Mode Decompositions (EMDs) and other complexity measures (such as entropy and fractal analysis). These features were calculated across all channels and a final median taken across channels as input into the RF model. The RF is an ensemble method that uses a large set (or ‘forest’) of trained decision trees to provide an averaged final prediction. Each tree is trained on a bootstrapped sample of the dataset and a random selection of the features which is shown to provide better prediction accuracy than from an individual tree and can also provide an implicit measure of the important features used in the algorithm. The RF model uses 1500 trees and utilizes all features for each tree split. All steps and hyperparameter choices used here are the same as our previous published RF approach (Pillay et al., 2020).

### 2.4 Model training and relative performance assessments using dataset 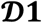

#### 2.4.1. *Splitting dataset* 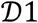 *into training set and test set*

Dataset 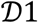 was used to train and test all models. By using a cohort of only normal outcome data, it is assumed that predicted brain age equates to true PMA. This allows training of a normative model to predict the PMA, and therefore brain age, for a normally developing baby (Pillay et al., 2020; Stevenson et al., 2020a, 2017). Further data can then be assessed against this trained model to identify deviations. Dataset 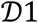 was divided by recording into age-stratified training and test sets of size 50 and 47 recordings, respectively (Supplementary Information S.2.; Supplementary Figure 1).

#### 2.4.2. *Model training in dataset* 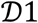

For the training of both deep learning models and the RF model, the mean squared error (MSE) loss was used. For the RF model, the model was re-trained using the original training procedure as previously outlined (Pillay et al., 2020). For the deep learning models, model training included early stopping, Gaussian noise addition, recording segmentation into 30 s segments, and ensemble learning. These four components are described in detail in Supplementary Information S.3, with early stopping, Gaussian noise addition, and ensemble learning included to increase robustness of the model.

#### 2.4.3. *Model testing in dataset* 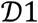

##### 2.4.3.1. Assessing model performance

The ultimate goal of each brain age prediction model is to generate a single brain age prediction estimate per EEG recording. For the deep learning models, each deep learning model generates ten brain age prediction estimates per 30 s segment of an EEG recording (as a 10-learner ensemble method was used, see Supplementary Information S.3.). During testing, all contiguous 30 s segments across each recording are used with the number of 30 s segments therefore dependent on the overall EEG recording duration. To aggregate a deep learning model’s predictions to a single value per recording, the median across the ten ensemble predictions per 30 s segment is determined, then a further median across all 30 s segments in the recording is taken resulting in the final prediction estimate. This is different to the RF model strategy, where a single brain age prediction estimate is generated per recording by manually calculating features in the 30 s segments and taking the medians across all segments before brain age prediction is performed. Across all recordings in the test set in 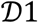, there were a total of 30K segments used. For both the deep learning models and RF model, the final prediction estimate for a recording is used to generate the prediction error (or absolute prediction error) for that recording.

##### 2.4.3.2. Reducing EEG channel requirements

The established RF model uses eight channels in a referential montage (Figure 1) to predict infant brain age. The performance of the RF model, the Sinc model, and the other deep learning models were assessed and compared using this initial setup. Subsequently, the deep learning models were re-trained and performances compared as the number of EEG channels were systematically reduced: 4-channel referential (C3, C4, T3 and T4), 2-channel referential (C3 and C4), and finally a 1-channel bipolar (C3-C4) montage (Figure 1). Channels were selected to ensure good symmetry across the midline of the scalp and ample coverage. The 1-channel bipolar montage was selected for its similarity to setups used in clinical amplitude-integrated EEG (aEEG) monitors. EEG pre-processing was independently repeated each time, with the amplitude standardisation step recalculated on the reduced channel configurations. After (re)training using the training set, each model generated a brain age prediction per recording in the test set. This set of predictions was used to generate a set of absolute errors per model. Using one-sample paired t-tests (p<0.05 significance level), we assessed the model performances by comparing t-statistic magnitudes and tested for statistically significant differences between the Sinc model’s mean absolute error and the mean absolute error of each of the alternative models (RF model and other deep learning models).

It is worth noting that the 1-channel bipolar montage used for our analyses was achieved by ignoring the additional channels unnecessary for this montage. This approach is distinct to a true clinical scenario when only a 1-channel bipolar montage would be used during recording. Our assumption, which we believe to be reasonable, is that both approaches to 1-channel bipolar montage setup are closely matched for this specific use case. However, this assumption should be tested in future external validations of the deep learning model using clinical grade bipolar montage data.

##### 2.4.3.3. Reducing EEG recording duration requirements

Having demonstrated the high performance of the deep learning Sinc model using the full-length EEG recording duration with only a 1-channel bipolar setup (section 2.4.3.2.), we next assessed the Sinc model performance using the 1-channel bipolar setup (Figure 1) as the EEG recording duration was systematically varied. We examined a range of recording lengths from 0.5–120 min and compared Sinc model performance on these reduced recording durations relative to the Sinc model performance with the full-length EEG recording duration to identify an appropriate reduced recording duration. To get a reduced recording from a single full recording, we randomly sampled each reduced duration segment from the full recording, generating an absolute error value per reduced duration segment. Due to the arbitrary nature of selecting a reduced recording segment from a full recording, we repeated the procedure using 1000 bootstrapped samples from which a mean absolute error was derived per recording per reduced recording duration. A minimum reduced recording duration was identified as the duration at which the prediction performance, measured using the mean absolute error, noticeably drops below that of the full duration Sinc model. Finally, the mean absolute error of this reduced duration (20 mins – see Section 3.1.) 1-channel Sinc model was compared to the mean absolute error of the 8-channel full duration RF model using a one-sample paired t-tests (p<0.05 significance level). For the reduced duration Sinc data, the initial 20 mins of each recording was selected for t-test analysis.

### 2.5. Interpreting Sinc model performance using dataset 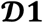

Deep neural networks are notorious for being black-box machines, limiting interpretability when compared to machine learning approaches and traditional visual assessment approaches. Nevertheless, methods are improving to visualize these networks to understand how they were trained and their potential to generalise well on new data. In this study, two visualization techniques were applied to further understand Sinc model performance: input-loss minimisation and uniform manifold approximation and projection, see Supplementary Information S.4.

### 2.6. External validation of Sinc model performance using dataset 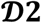

The final Sinc model was trained on the entire dataset 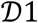 using the 1-channel bipolar setup (Figure 1). Similarly, the final “gold standard” RF model was trained on the entire dataset 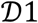 using the 8-channel referential setup. When applying the final Sinc model to the independent hold-out dataset 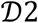, the 1-channel bipolar setup was used and a 20 mins recording duration was randomly sampled from the full duration EEG recording. When applying the final RF model to dataset 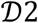, the 8-channel referential setup and full EEG recording were used.

#### 2.6.1. PMA prediction in independent hold-out dataset

To assess the generalisability of the Sinc model to predict infants’ PMA on independent data, the normal BSID-II outcome data from the independent hold-out dataset 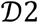 was used. To assess the association between true PMA and predicted PMA, a linear mixed effects regression model was used (p<0.05 significance level). Random intercepts were introduced to group repeated recordings from the same infant. Associations between true PMA and predicted PMA were also assessed for the mild abnormal and severe abnormal groups. Similarly, the RF model was used to generate PMA predictions for the normal, mild abnormal, and severe abnormal groups in dataset 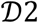, and associations between true age and predicted age were assessed in an identical manner to the Sinc model.

The two models’ PMA prediction performances were compared using the linear mixed effects regression models’ z-statistic magnitudes per BSID-II outcome cohort. Additionally, Bland-Altman analysis (Bland and Altman, 1999, 1986) was used to assess Sinc-RF model agreement in absolute error magnitude, pooled across all data in dataset 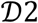. Bland-Altman analysis was implemented in R v4.1.1 (R Core Team, 2018) using a publicly available package (https://rdrr.io/cran/BlandAltmanLeh) to estimate the bias (Sinc minus RF) and limits of agreement, along with 95% confidence intervals. Model agreement was assessed using individual recording absolute errors and using within-infant multi-recording mean absolute errors, to assess the influence of within-infant multi-recording averaging on model agreement (Bland-Altman plot y-axis, limits of agreement width) and average prediction error magnitude (Bland-Altman plot x-axis, range).

#### 2.6.2. Associating brain age delta magnitude to 9-month BSID-II follow-up outcomes

The association between an infant’s brain age delta magnitude and 9-month BSID-II follow-up outcomes (normal, mild abnormal, severe abnormal) was assessed for all infants in dataset 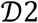. For each infant, a brain age delta (absolute error) was determined per recording, and the mean absolute error (MAE) across an infant’s multiple recordings was used as an estimate of that infant’s brain age delta i.e. the deviation between their brain age and their true age. This per-infant MAE thus represents an infant’s overall brain neurodevelopmental trajectory deviation, with a larger trajectory deviation corresponding to greater deviations from the norm.

Trajectory deviations across all infants in dataset 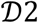 were then grouped by neurodevelopmental outcome (as defined in section 2.1.1.) and significant differences between groups assessed using one-way ANOVA (p<0.05 significance level). Tukey’s post-hoc test, which corrects for multiple comparisons (p<0.05 significance level), was used to identify significant pair-wise comparisons. Additionally, the two models’ BSID-II outcome group separation performances were compared using the pairwise standardised effect size (Cohen’s D, estimated using MATLAB’s meanEffectSize function) magnitudes per contrast: mild minus normal, severe minus mild, and severe minus normal.

Finally, group-wise (normal, mild abnormal, severe abnormal) differences in GA, PMA and the number of recordings in each infant’s trajectory were checked using one-way ANOVA to assess their potential influence as confounding factors.

### 2.7. External validation of Sinc model performance using dataset 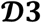

The final Sinc model was applied to the independent dataset 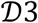 collected at an independent centre (Oxford, UK). The 1-channel bipolar montage (C3-C4) and the first 20-minutes of each recording were used in the analysis.

The association between true PMA and predicted PMA was assessed using Pearson correlation (z-statistic calculated using the Fisher r-to-z transform, p<0.05 significance level). Z-statistics are reported for the results of both datasets 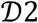 and 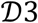. Each infant in dataset 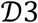 was recorded on a single test occasion; the group-level MAE was calculated as the mean across all recordings of each subject-level brain age delta i.e. each infant’s error in predicted versus true age.

The brain age delta estimate can have a dependency with age – an age association bias that is known to occur for several distinct reasons such as regression dilution (Smith et al., 2019). To correct for this age association bias, we adjusted the predicted brain age using the linear regression between the brain age delta and the true age (Smith et al., 2019). To assess the generalisability of this correction to new data we adjusted the predicted brain age using leave-one-subject-out cross-validation, calculating the MAE of the held-out subject compared with its true age.

## 3. Results

### 3.1. The Sinc model outperforms alternative model architectures in predicting infant age (dataset 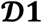)

A comparison of model performance across the Sinc model and four alternative candidate deep learning models, with reduced channel setups is summarised in Figure 3a and Supplementary Table 1. Using the 8-channel setup and the full recording duration data of dataset 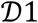, the Sinc model out-performed both the established benchmark RF model (Sinc error = 0.73 weeks, RF error = 1.01 weeks, n = 47 recordings, t-statistic = 1.44, p = 0.078) as well as the candidate deep learning models. When the number of recording channels was reduced from eight to one (bipolar channel, C3-C4), the Sinc model had consistently lower MAE values compared with alternative models and exhibited a total drop in performance of only 0.05 weeks (Sinc: 8-channel MAE = 0.73 weeks, 1-channel MAE = 0.78 weeks). Furthermore, the 1-channel bipolar Sinc model outperformed the 8-channel referential RF model (Sinc error = 0.78 weeks, RF error = 1.01 weeks, n = 47 recordings, t-statistic = 1.13, p = 0.13).

**Figure 3:**
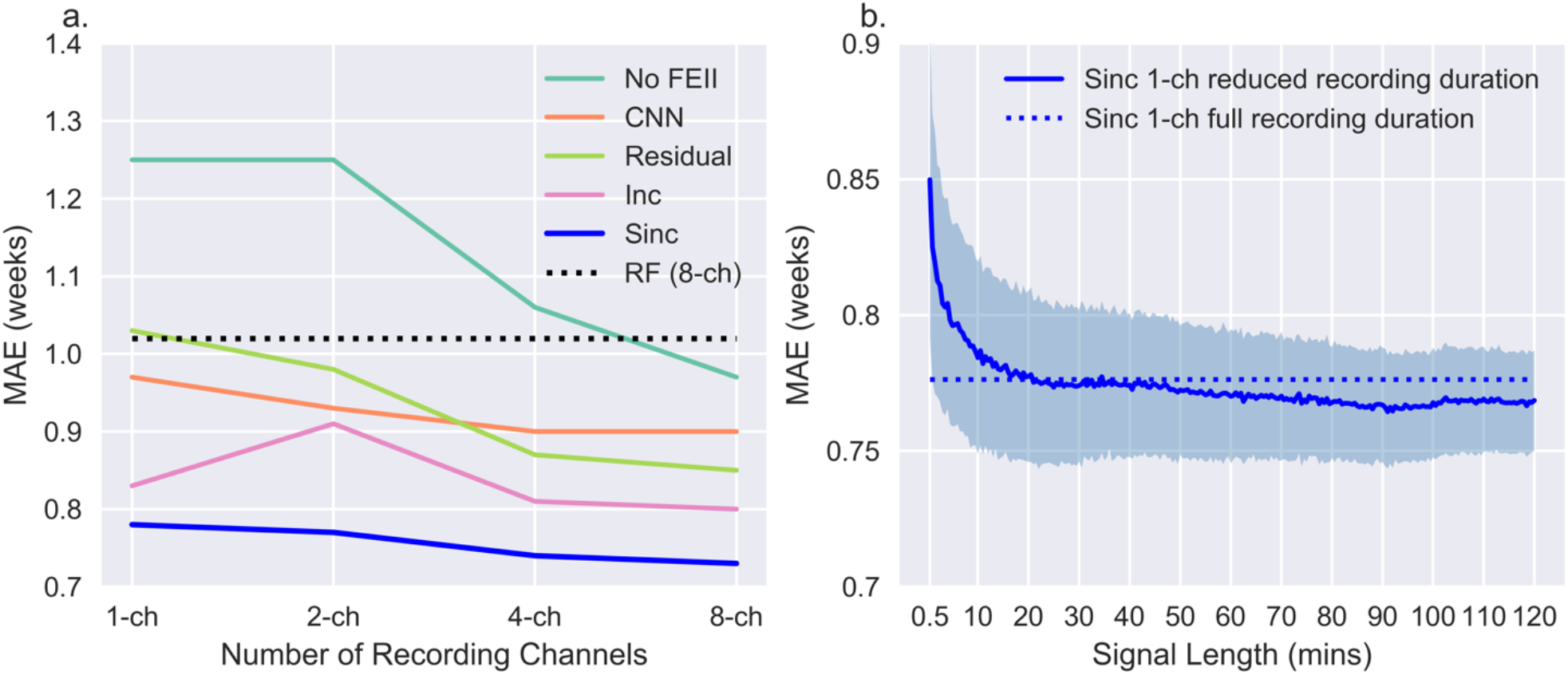
The Sinc model outperforms alternative architectures in predicting infant brain age. Brain age prediction performance (MAE) using dataset 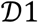 test set. **a.** Each line represents a different model, and each model uses the entire recording duration. See Supplementary Table 1 for plotted values. The RF model is the established benchmark, which uses eight channels. The Sinc model consistently outperforms both the RF model and the alternative deep learning models, with a lower prediction error using a single channel (MAE = 0.78 weeks) than the RF model using eight channels (MAE = 1.01 weeks). **b.** The Sinc model’s performance using a single channel and the full recording duration (MAE = 0.78 weeks, dotted line) was used as a benchmark to assess Sinc model performance with a single channel and systematically reduced recording durations (solid line). Performance using the reduced recording durations are matched to the full recording duration when recordings of 20 mins or longer are used; using less than 20 mins recording duration exhibits a gradual drop in prediction performance. Shaded intervals denote the standard deviation for the reduced recording durations. Note, MAE performance suggests a drop below the full signal performance beyond 20 min duration. This is due to the bootstrap sampling error (Efron and Tibshirani, 1994), and this inherent bias is a fluctuation about the full recording MAE with standard deviation <1. We can assume that the MAE beyond 20 mins is equivalent to the MAE when the full recording duration is used. As it is too computationally intensive to show performance beyond 2 hour signal durations the random variation cannot be fully shown here. Abbreviations: FE = feature extraction; CNN = convolutional neural network; Inc = inception; Sinc = shared inception; RF = random forest; ch = channel; MAE = mean absolute error.

The Sinc model prediction error recorded from a single channel with full recording duration (duration: median = 4h 25m, IQR = 4h 4m–7h 10m) was compared to Sinc model prediction error using a single channel and reduced recording durations ranging from 0.5–120 mins (Figure 3b). Using only 20 mins of EEG recording, the mean Sinc model prediction error was equivalent to using the full recording duration. Using the established RF method as a benchmark, which relied on an 8-channel setup and full-length recordings, the proposed Sinc model outperformed this benchmark while having practical setup requirements that are far more achievable and practical for use in a clinical environment (Sinc error = 0.79 weeks, RF error = 1.01 weeks, n = 47 recordings, t-statistic = 1.07, p = 0.14). While the three Sinc models’ performances (8-channel full duration, 1-channel full duration, 1-channel 20 min duration) did not statistically significantly differ to the benchmark RF model performance, the Sinc model’s performances were marginally but consistently improved (t-statistics = 1.44, 1.13, 1.07, respectively, with positive t-statistics indicating larger MAE for RF).

### 3.2. Sinc model may determine age using degree of EEG continuity (dataset 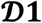)

To shed light on the specific EEG features that the deep learning Sinc model is likely utilising for the brain age prediction, a method called input-loss minimisation was used to generate synthetic EEG data that would force the model to make a brain age prediction of 30 weeks, 35 weeks, and 40 weeks PMA, respectively (Figure 4). Visually examining the synthetic EEG data shows that EEG continuity and bursting were qualitatively distinguishing features and are therefore likely features that the Sinc model used to characterise age-dependent activity. The 30-week synthetic data reflects aspects of high discontinuity with short, high amplitude bursts and long-duration inter-burst intervals (approximately 5-20 s) (Figure 4a). With increasing PMA, the inter-burst interval durations decreased and burst periods widened, and by term-age, the signal was almost fully continuous with no clear burst or inter-burst interval patterns (Figure 4c).

**Figure 4:**
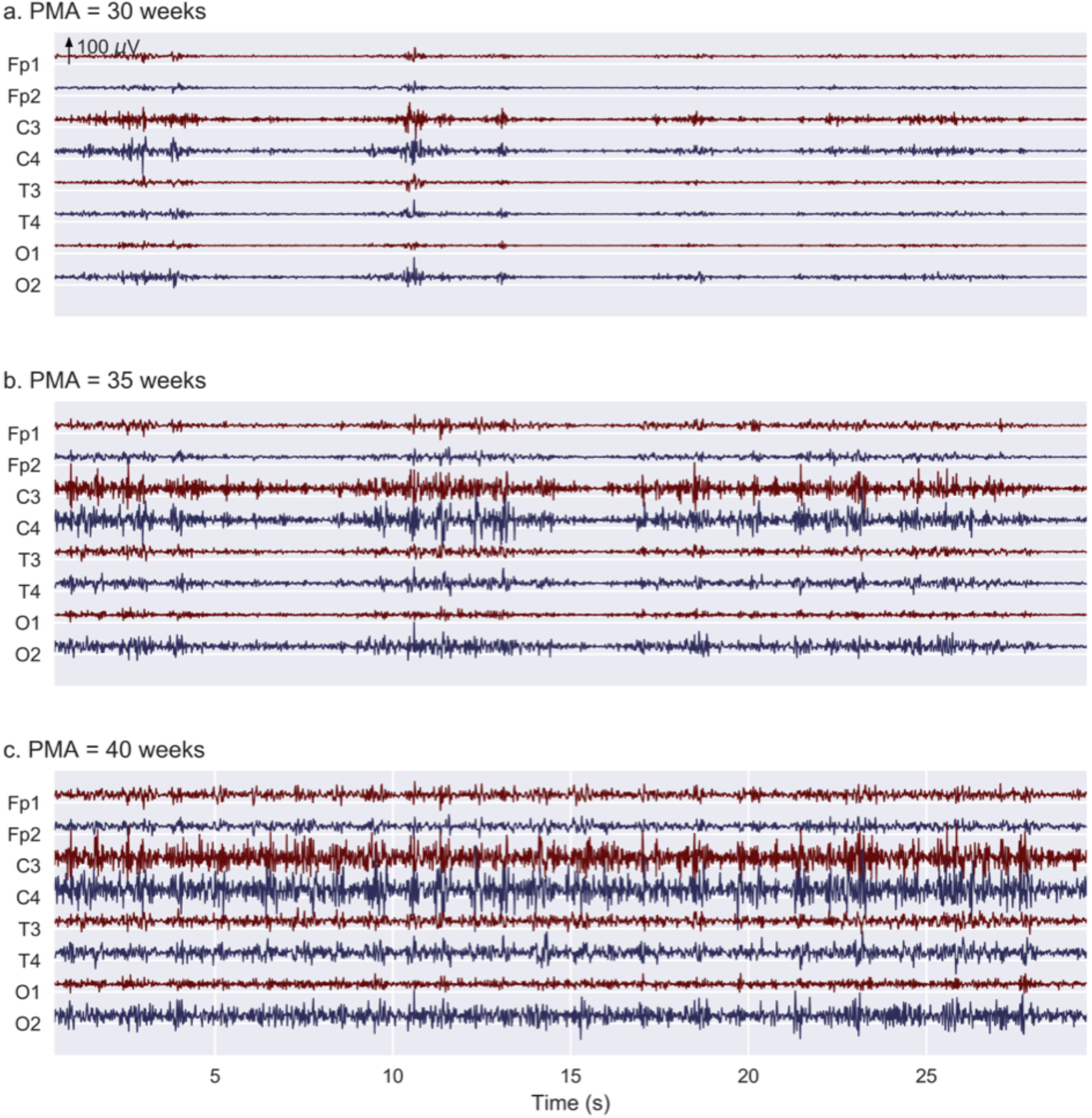
Synthetic EEG data generated using the Sinc model highlight changes in discontinuity characteristics with PMA, reminiscent of maturational trends seen in real EEG data. Results are generated using the input-loss minimization technique for three target PMAs (30, 35, and 40 weeks) spanning the early preterm to term age range. This is performed for the 8-channel full recording duration case. The degree of continuity in activity can be seen to increase with PMA.

Using UMAP to visualise the data inputs to the three Sinc blocks (FEI, FEII, and Regression), a clear separation of features occurs, beginning with a low-level followed by high-level feature extraction (Figure 5). At the stage of inputs to Regression, the data can visually be seen to separate such that datapoints increase almost monotonically with PMA (Figure 5c). This clear progression is indicative that the network weights are trained well in the intermediate layers, and this visualisation provides further insight into the role of each block.

**Figure 5:**
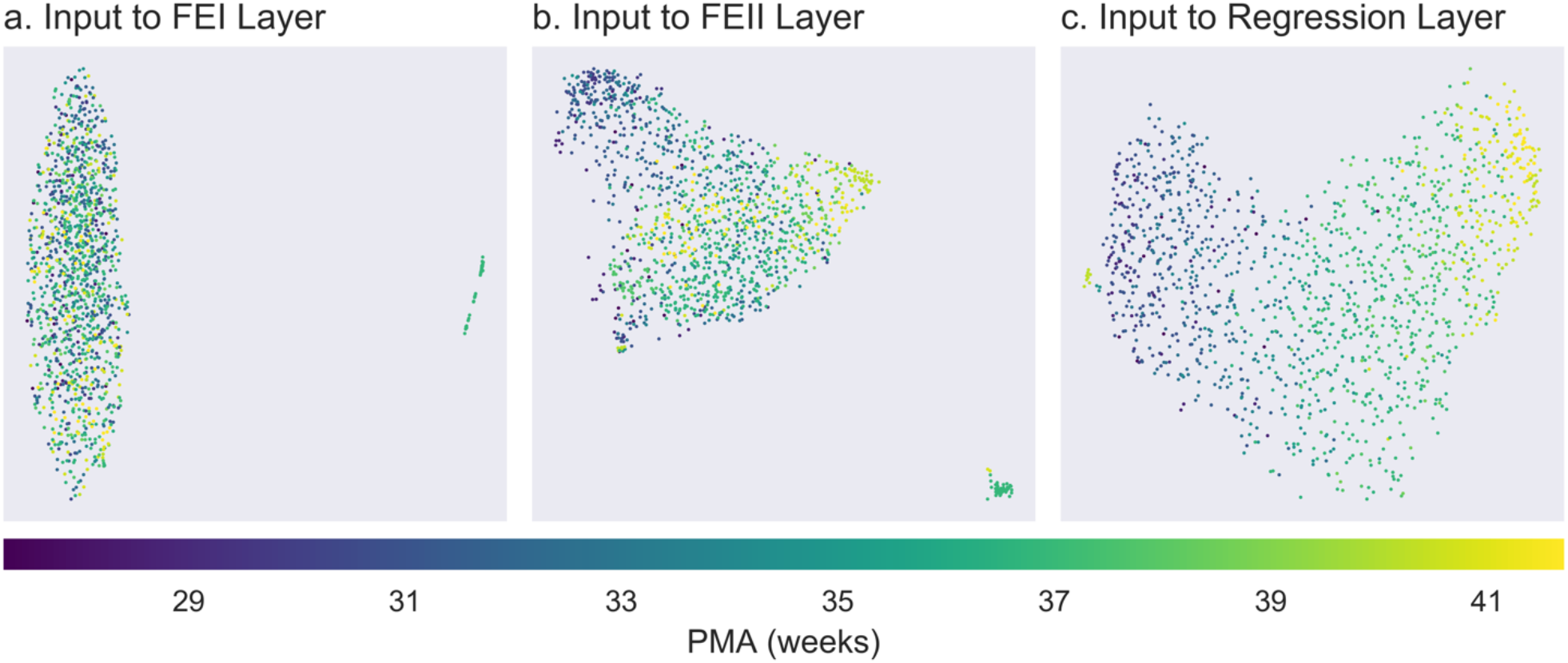
Visualising Sinc model performance using UMAPs. Visualization of the inputs at various blocks in the proposed model: Feature Extraction I (FEI), Feature Extraction II (FEII), and Regression (see Figure 1b). Results are shown for the 1-channel, full recording duration case in Dataset 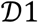. An increasing separation of the features with respect to PMA is seen on moving from a-c. This clear progression is indicative that the network weights are trained well in the intermediate layers, and this visualisation provides further insight into the role of each CNN block. **a.** Input to FEI has not yet been processed, so there is no separation of inputs to FEI. **b.** The input to FEII is the output from FEI. It is evident that the role of FEI is to perform a low-level ‘feature extraction’ that performs an initial separation between the very preterm (blue dots) and preterm and term age groups (green and yellow dots) i.e. a general separation between strong discontinuity and continuity in the EEG. **c.** The input to Regression is the output of FEII. The FEII stage performs a higher-level feature extraction providing further discriminatory power, allowing better separation of these mid-age (31-37 weeks) and term age groups. Furthermore, at the stage of input to Regression, we observe that the PMAs of the datapoints from left to right increase almost monotonically such that the very left and right datapoints correspond to the extremely young and old neonates, respectively, while the middle ages are almost uniformly distributed in-between. Abbreviations: UMAP = uniform manifold approximation and projection.

### 3.3. Sinc model brain age prediction generalises accurately to an independent hold-out dataset (dataset 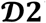)

Having established the Sinc model in dataset 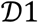 (section 3.1), this model was applied to a healthy cohort of infants’ data from the independent hold-out dataset 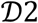. Using 1-channel bipolar EEG data of 20 min recording duration, the Sinc model’s predicted ages were statistically significantly correlated with infants’ true PMA (Normal: n = 22 infants, z-statistic = 33.32, p < 0.0001) (Figure 6ai), demonstrating that the model successfully generalises to independent data. The Sinc model also generated predicted ages that were statistically significantly correlated with infants’ true PMA for the infants in dataset 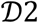 that had abnormal BSID-II follow-up outcomes (Mild abnormal: n = 10 infants, z-statistic = 18.03, p < 0.0001; Severe abnormal: n = 10 infants, z-statistic = 15.54, p < 0.0001) (Figure 6aii). Infants with abnormal BSID-II follow-up outcomes were not used in training the Sinc model, and so age predictions for these cohorts were, as expected, less accurate than those of the healthy outcome cohort and thus exhibited weaker correlations (although still very strong) between brain age and true age.

**Figure 6:**
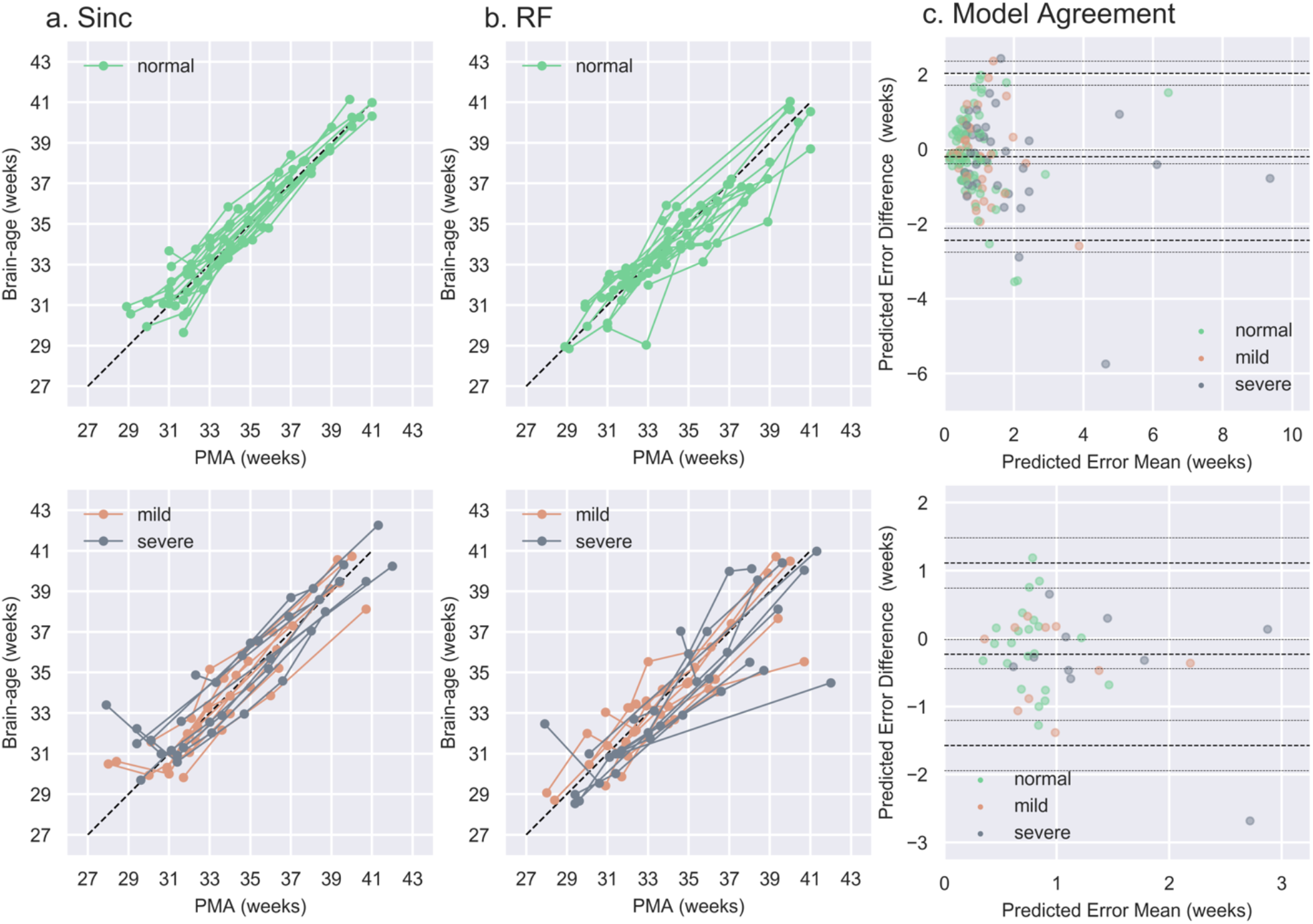
Brain age prediction models generalise to independent dataset 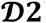. **a.** Sinc model brain age predictions for infants with (i) normal and (ii) abnormal BSID-II follow-up outcomes (Dataset 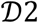). Each string of connected points is a single infant’s longitudinally-assessed multi-recording trajectory, and the dashed black line is the y=x line along which perfect predictions would lie. **b.** RF model brain age predictions for infants with (i) normal and (ii) abnormal BSID-II follow-up outcomes. **c.** Bland-Altman plots to assess agreement between Sinc and RF models’ PMA prediction performances, quantified using absolute prediction errors. In both plots, the x-axis is the mean prediction error of the two models, and the y-axis is the difference in prediction errors (Sinc minus RF). The heavy grey lines are the mean bias and limits of agreement, while the light grey lines indicate the 95% CI for the bias and limits of agreement. (i) Per-recording model agreement assessment. (ii) Per-infant model agreement assessment i.e. multi-recording average per infant. Note the greater model agreement (narrower limits of agreement along y-axis) and reduced average prediction error (shorter range along x-axis) when using the multi-recording average prediction error in (ii) compared to the single recording prediction error in (i). Abbreviations: Sinc = shared inception; RF = random forest; PMA = postmenstrual age.

Using the 8-channel EEG setup and the entire recording duration, the RF model generated age predictions that were statistically significantly correlated with infants’ true PMA for both the normal outcome and abnormal outcome cohorts (Normal: z-statistic = 22.89, p < 0.0001; Mild abnormal: z-statistic = 12.51, p < 0.0001; Severe abnormal: z-statistic = 10.76, p < 0.0001) (Figure 6b). While the brain age prediction correlation results for both the novel Sinc model and the established RF model were very strong and highly significant for all three infant cohorts, the Sinc model consistently outperformed the RF model per cohort (consistently larger z-statistics). Importantly, Sinc’s improved prediction accuracy was achieved while using dramatically lower EEG data requirements.

To quantitatively assess the level of agreement in PMA prediction performance between the RF and Sinc models, we generated Bland-Altman plots of absolute prediction errors for the entirety of dataset 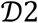 (pooled normal, mild abnormal, and severe abnormal outcome data) based on both individual recordings (n = 141 recordings in total) (Figure 6ci) and individual infants (n = 42 infants in total) (Figure 6cii). In both instances, there was a statistically significant negative bias reflecting the reduced prediction error using the Sinc model (per-recording: mean bias = −0.202, 95% CI = [−0.387, −0.016]; per-infant: mean bias = −0.231, 95% CI = [−0.444, −0.017]). Assessing the individual recordings data, the limits of agreement were −2.435 and 2.032 with 95% CI = [−2.756, −2.115] and [1.712, 2.353], respectively (Figure 6ci). Assessing the individual infants’ data (multi-recording average per infant), the limits of agreement were −1.573 and 1.112 with 95% CI = [−1.943, −1.204] and [0.742, 1.481], respectively (Figure 6cii). The narrower limits of agreement width using the infant-level assessment highlights a noticeable increase in Sinc-RF model agreement when using multi-recording average prediction errors per infant rather than prediction errors based on individual recordings, due to the reduced random noise variance as raeconrdseinqguaevnecreagoifngth. eUsminugltmi-ulti-recording average prediction errors per infant, we can expect 95% of absolute prediction error differences between the RF and Sinc models to be approximately ±1.5 weeks, and the Sinc model to have a smaller prediction error of approximately 0.23 weeks on average.

### 3.4. Sinc model brain age deltas are associated with 9-month follow-up neurodevelopmental outcomes (dataset 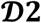)

The variability in brain age delta magnitudes between infants with normal and abnormal BSID-II follow-up outcomes forms the foundation of the possibility of using brain age prediction to risk-stratify infants in the first few weeks of postnatal life according to neurodevelopmental outcomes. Here, using the Sinc model, the average brain age deltas for the normal, mild abnormal, and severe abnormal outcomes groups assessed using the BSID-II at nine months postnatal age were found to significantly differ (Normal: mean MAE = 0.71, n = 22 infants; Mild abnormal: mean MAE = 0.79, n = 10 infants; Severe abnormal: mean MAE = 1.27, n = 10 infants; one-way ANOVA: f-statistic = 4.24, p = 0.02) (Figure 7a). Significant differences between the mean deltas for the normal and severe abnormal groups were observed using post-hoc analysis adjusted for multiple comparisons (Tukey test: q-statistic = 4.20, p = 0.02) (Figure 7a). Taken together, these results indicate that Sinc model brain age delta magnitudes, generated using a single channel and 20 mins recording duration, scale with clinically informative BSID-II outcomes that are assessed several months later.

**Figure 7:**
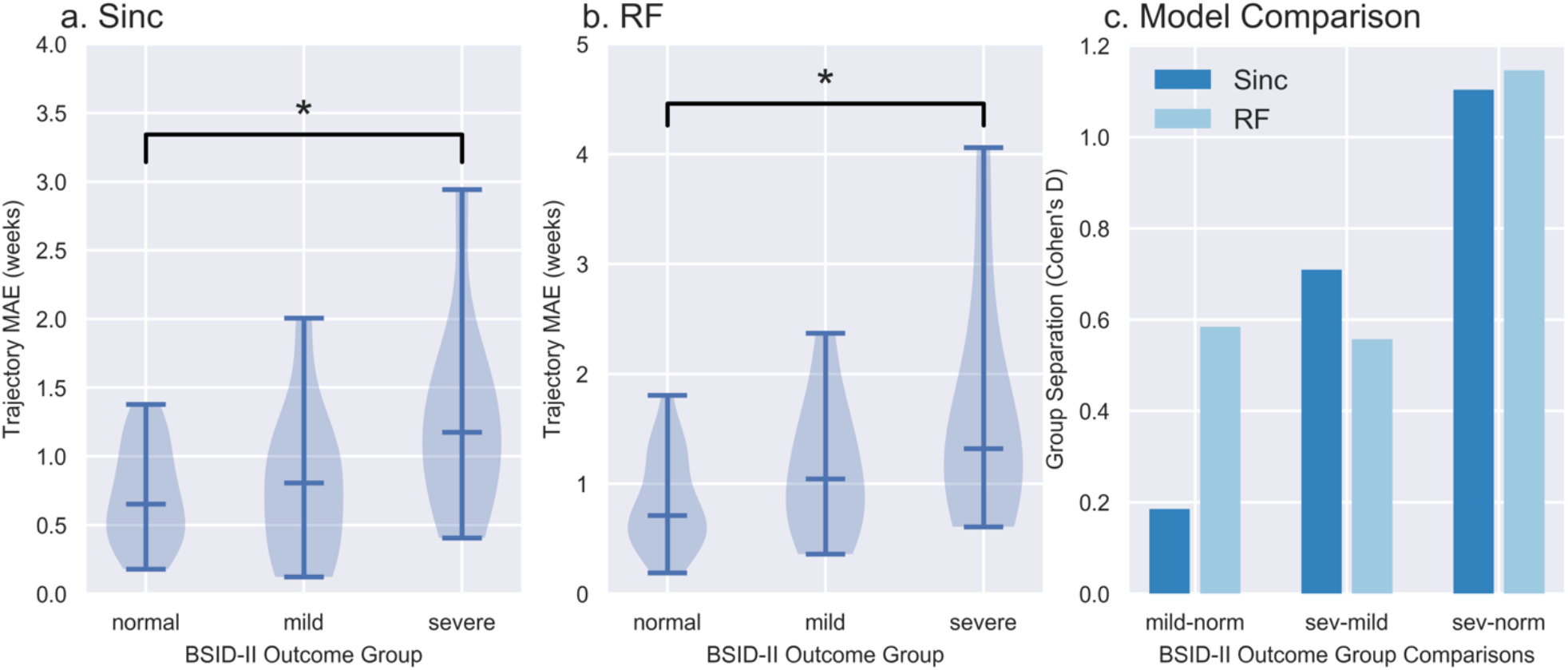
Brain age delta magnitudes scale with 9-month follow-up neurodevelopmental outcomes. **a.** Sinc model absolute prediction error magnitudes (brain age deltas) for each of the three BSID-II outcome cohorts: normal, mild abnormal, and severe abnormal. The average prediction error is larger for poorer 9-month follow-up BSID-II neurodevelopmental outcomes, and the mean prediction error for the severe abnormal group is significantly larger than that of the normal group. **b.** RF model absolute prediction error magnitudes for each of the three BSID-II outcome cohorts. The average prediction error is larger for poorer 9-month follow-up BSID-II neurodevelopmental outcomes, and the mean prediction error for the severe abnormal group is significantly larger than that of the normal group. **c.** The x-axis displays each of the three combinations of pairwise comparisons for the three BSID-II outcome cohorts: mild minus normal, severe minus mild, and severe minus normal. For each model, the y-axis displays the standardised effect size (Cohen’s D) separating each pair of BSID-II outcome cohort. Sinc = shared inception; RF = random forest; MAE = mean absolute error; BSID-II = Bayley scale of infant development; * = statistically significant.

As reported previously, the RF model’s brain age deltas also significantly differed between the three BSID-II outcome cohorts (Normal: mean MAE = 0.83, Mild abnormal: mean MAE = 1.13, Severe abnormal: mean MAE = 1.63, one-way ANOVA: f-statistic = 4.96, p = 0.01) (Figure 7b), with significant differences observed between the mean prediction errors for the normal and severe abnormal groups (Tukey test: q-statistic = 4.36, p = 0.01) (Figure 7b).

Quantitatively assessing the magnitude of the group average MAE separation between BSID-II outcome cohorts, a similar trend was observed for both the Sinc and RF models (Figure 7c). Both models exhibited poorest separation between the normal and mild abnormal outcome cohorts (group separation effect size: Sinc Cohen’s D = 0.186; RF Cohen’s D = 0.585), an intermediate degree of separation between the mild abnormal and severe abnormal outcome cohorts (group separation effect size: Sinc Cohen’s D = 0.71; RF Cohen’s D = 0.557), and greatest separation between the normal and severe abnormal outcome cohorts (group separation effect size: Sinc Cohen’s D = 1.104; RF Cohen’s D = 1.146) (Figure 7c).

No significant differences were identified between outcome groups for the potential confounding variables. Sinc model MAEs one-way ANOVA results (n = 42): GA: f-statistic = 0.93, p = 0.40; PMA: f-statistic = 0.51, p = 0.60; trajectory recording number: f-statistic = 0.28, p = 0.76).

### 3.5 Sinc model accurately predicts brain age in data collected at an independent site (dataset 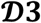)

The Sinc model was applied to an independent dataset collected at an independent centre (Oxford, UK; dataset 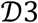). The Sinc model’s predicted ages were significantly correlated with the infant’s true PMA (n = 73 infants, Pearson correlation coefficient r=0.91, z-statistic=1.52, p < 0.0001, Figure 8a), with good prediction accuracy (MAE = 0.97 weeks). This highlights that the Sinc model can generate age predictions using single recordings per infant for accurate group-level analysis at an independent site.

**Figure 8:**
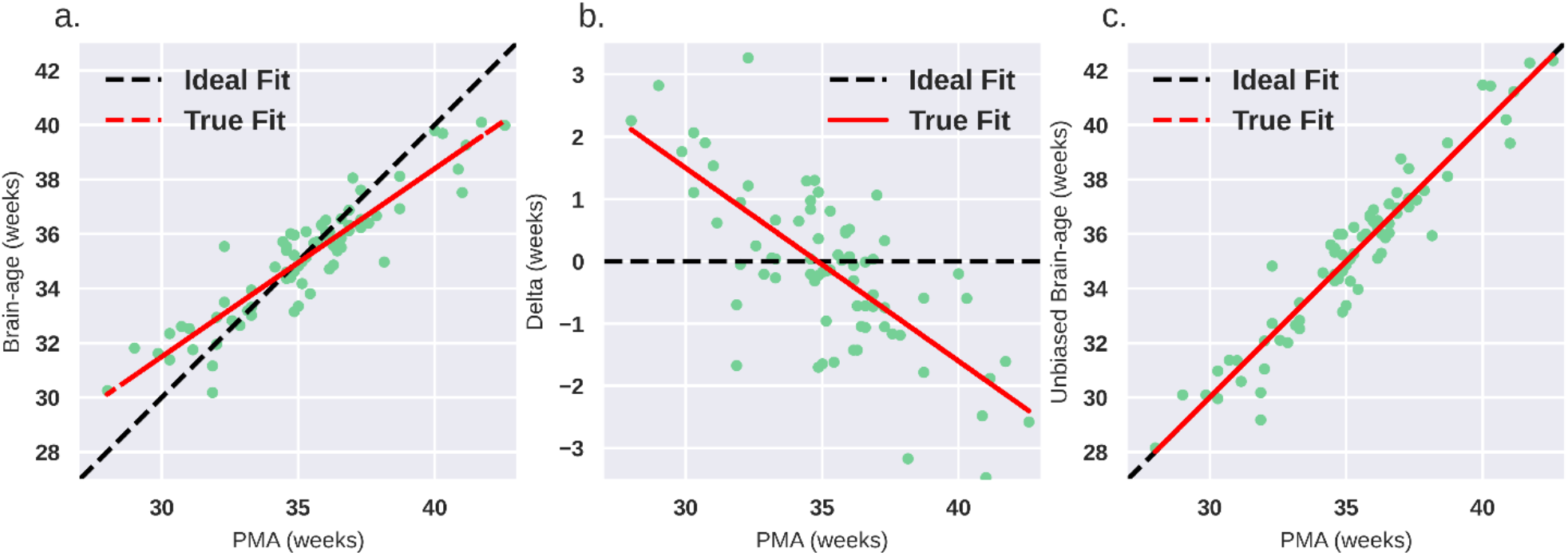
Sinc model brain age prediction generalises to dataset 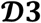. In each panel (a-c), each point indicates a single infant (n=73); the dashed black line is the ideal fit line; and the red solid line is the true fit line (least squares). **a.** Sinc model brain age predictions for dataset 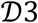. The ideal fit line is the y=x line of perfect prediction. The misalignment between the ideal fit line and the true fit line indicates an age association bias. **b.** Correlation between the brain age delta (predicted age minus true age) and the infant’s true age. The ideal fit line is the y=0 line of zero age association bias. The slope of the true fit line indicates the magnitude and direction of the age association bias. **c.** The predicted brain age after adjusting for the delta age association bias using leave-one-out cross validation. The ideal fit line is the y=x line of perfect prediction.

Unlike dataset *D2*, a noticeable bias in age prediction was visible in dataset 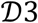 (Figure 8a). The magnitude of the brain age delta was significantly negatively correlated with the infant’s true PMA (r =-0.24, p<0.01, Figure 8b). To generate unbiased brain age delta values, this age association should be minimised (Smith et al., 2019). A simple linear regression model trained on dataset 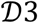, and validated using leave-one-out cross-validation, reduces this bias (Figure 8c). This additional linear model could be used in novel single-subject data collected at this site to produce brain age deltas with minimal age association bias. However, the biological value of the brain age deltas in dataset 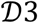 has yet to be established. This dataset currently does not have follow-up BSID-II outcomes, so the association between brain age deltas and follow-up outcomes could not be assessed.

## 4. Discussion

This study presents the first deep learning architecture for the prediction of brain age from infant EEG activity. The model is based on a deep CNN structure incorporating the new Sinc block for enhanced multi-scale decompositions, with prediction likely utilising between-infant differences in their EEG continuity and bursting characteristics. Relative to previous proof-of-concept studies (Pillay et al., 2020; Stevenson et al., 2020a), the current deep learning approach was able to predict infant brain age with comparable accuracy and generate brain age delta magnitudes that were significantly associated with neurodevelopmental outcome at a 9-month follow-up using BSID-II assessment. Importantly, the current approach achieved this using dramatically reduced EEG data utilisation requirements, relying on only a single channel bipolar montage and 20 mins recording duration. This is important as it suggests that future systems utilising this method may only require single-channel capabilities which is simpler to set up and makes EEG data acquisition easier. This streamlined model, which can be applied in an objective and automated manner, thus demonstrates potential clinical utility for cot-side monitoring assessment of neurological well-being.

The chosen development strategy for the Sinc model involves training and testing the model first on a normal development dataset 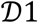 and then additionally assessing performance in two independent datasets (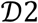 and 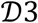, the latter collected at an independent site). Although we performed a single split on 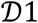 for initial training and testing and could have used alternative techniques (such as cross validation), the goal was to assess relative performance with this dataset when comparing models, channel numbers, and recording durations. We kept the training and test splits in 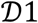 consistent across these comparisons ensuring that relative differences in performance were meaningfully comparable. Furthermore, by showing high performance in the brain age prediction in the independent datasets, which was comparable to the held-out test set performance in 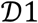, we can justify with confidence that the training strategies and choices made have still resulted in a robust generalisable model.

The model performed well on data collected at an independent site, despite differences in data collection such as EEG recording equipment and research personnel. This importantly suggests that the model is generalisable and could easily be employed for clinical use across multiple hospitals. Interestingly, an age association bias in model estimates could be observed between the predicted age and true age when the model was applied to dataset 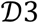 (Oxford dataset), with the model to overestimate age in the youngest infants and underestimate age in the oldest infants. The bias was not observed in dataset 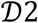 (Leuven dataset). Bias in brain age predictions can arise from number of factors (Smith et al., 2019): for example, “regression dilution” due to errors in measurement of the predictors (dataset 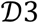 used single recordings per infant, while dataset 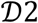 used multiple recordings per infant affording reduced measurement error). Using leave-one-subject-out cross-validation, we demonstrated that it was possible to minimise this bias in dataset 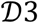, suggesting that this correction would be generalisable for future infants collected at this centre.

Throughout our analyses, we used our previously published (Pillay et al., 2020) RF model as a “gold standard” benchmark against which our novel Sinc model’s performance was assessed. The RF model used an 8-channel referential montage, over an hour of EEG recording, required sleep-staging and an explicit pre-calculation of over 200 established features, while the Sinc model required only a 1-channel bipolar montage and a 20 min recording duration, no sleep-staging, and included an implicit feature extraction step. In all analyses, the Sinc model either performed comparably to or out-performed the RF model. Additionally, in work published by an independent group (Stevenson et al., 2020a), brain age deltas exhibited greatest separation between infants with normal and severely abnormal BSID-II follow-up outcomes – an observation that is consistent with the current study’s findings, further supporting the results of the Sinc model.

Although a quantitative analysis of model speed was beyond the scope of this study, it is clear from previous studies (Pillay et al., 2020; Stevenson et al., 2020a) that the requirement to extract multiple features (some highly complex and non-linear), as well as the need to pre-stage the EEG based on sleep state or states of discontinuity would slow performance, and this is suggested in a related study on neonatal sleep-staging (Ansari et al., 2018). With the right accelerated hardware, however, the proposed model (once trained) performs brain age predictions very quickly. This simplified analysis pipeline lends itself well for hospital use if fast feedback is required in high-intensity contexts, for instance, while the infant is in critical or post-operative care.

A further advantage of the Sinc model over the other deep learning architectures tested here is the introduction of the Sinc block which, with a reasonable number of parameters, achieves a highly non-linear architecture for performing multi-scale analysis (Ansari et al., 2021). The streamlined preprocessing and feature extraction as well as the highly non-linear nature of the Sinc model are invaluable attributes that provide flexibility for extraction of key signal characteristics and result in a more focused feature set. The deep learning Sinc model is thus a flexible and efficient approach for use with neonatal EEG data, which are data that typically exhibits highly variable and diverse signal patterns.

Using the trained Sinc model to generate synthetic EEG data (Figure 4), our results suggest the model’s predictive performance may rely on identifying signal characteristics related to changes in the EEG discontinuity with age (related to bursts and inter-burst intervals). This finding relates sensibly to other findings in the current paper as well as established understanding of infant EEG maturation. Regarding our present findings, the Sinc model’s performance did not drop substantially going from eight channels to one, or full recording duration to 20 mins. This might suggest that the feature extraction stages of the architecture may be more tuned to global channel-independent characteristics (such as bursting and continuity), as opposed to spatially-dependent characteristics (such as inter-channel synchrony). Further, if the model relies on identifying changes in burst/inter-burst cycling and encodes this in a highly multi-scale manner, this may indicate that information on an infant’s burst/inter-burst cycling may be sufficiently discernible from a 20-minute EEG recording, with additional data providing diminished returns in discriminatory power.

Regarding infant EEG maturation, the progression of burst/inter-burst activity to continuous activity is the expected characteristic developmental trajectory from preterm to term age (André et al., 2010). Interestingly, these discontinuity patterns are also key for human experts when performing visual age prediction (Dereymaeker et al., 2017a; Husain, 2005). Observing this link between the synthetic inputs generated by the trained model and expected maturational trends strongly suggests the Sinc model is relying on biophysiologically sensible signal features, which is important for the generalisability of a model to novel data. We can tentatively suggest further similarities between the Sinc model’s generated synthetic EEG data and prominent features in the RF model. In agreement with our previous work (Pillay et al., 2020), prominent features chosen by the comparison RF model retrained in this study were based on the Line Length Burst %, a measure of the percentage of burst periods in the EEG (Koolen et al., 2014), as well as measures of skewness of the EEG amplitudes, which measure the asymmetry of a distribution compared to a Gaussian distribution. Line Length Burst % would be expected to change with PMA as the burst periods decrease with age and the EEG transitions to a more continuous pattern. Similarly, during this transition, the distribution shifts away from a symmetrical Gaussian distribution as the number of high positive bursts or spike amplitudes decreases. When comparing to the simulated results of Sinc in Figure 4, we see similar behaviour is also identified by this trained neural network emphasising the importance of this EEG characteristic across age.

We also note potentially interesting amplitude effects that are visible when looking at the model’s synthetic data across eight channels. For example, channels C3 and C4 have larger signal amplitudes relative to other channels. While amplitude is a feature that changes with maturation (André et al., 2010) making inter-subject variability in amplitude of potential value for brain age prediction, one must be cautious when interpreting this subtler cross-channel amplitude effect in the synthetic data. These amplitude effects may reflect a biophysiologically interesting phenomenon or may be an artefactual consequence of proximity to the Cz reference electrode. Future work on the Sinc model may help shed light on the potential role of amplitude effects.

Additionally, the role of motion artefacts, potentially related to sleep state and general motor activity levels, could influence prediction performance. We applied a very simple amplitude-threshold approach for artefact removal, and while this eliminates any major baseline drifts, periods of recording drop-off or high-amplitude motion artefacts, some subtler artefacts likely remain. It is unclear whether any residual motion effects influence prediction performance (either beneficially or detrimentally). However, the lack of motion-like signals in the model-generated synthetic EEG data suggests motion is unlikely to be playing a major role.

The ultimate interest in studying brain age delta magnitude is that neurological dysfunction can manifest in infants’ EEG as both accelerated or slowed maturation relative to a normative trajectory (Scher, 1997; Watanabe et al., 1999), and these functional maturational deviations have prognostic value (Iyer et al., 2015; Tokariev et al., 2019). The present study focused on the prognostic value of preterm and term age resting-state brain function as a basis for risk-stratification using 9-month BSID-II follow-up as the relevant outcome. However, as with any scale, there are limitations to BSID-II predictive validity (Hack et al., 2005). Clinical decision making regarding the provision of developmental care interventions (Burke, 2018) using deep learning-based predictions of infant brain age would benefit from advancing the prognostic validity of the brain age delta metric. For example, demonstrating associations between the metric and additional follow-up outcome metrics, such as executive function (Dai et al., 2021), would improve validity. Additionally, understanding the association between the metric and contemporaneous structural (e.g. body weight, brain structural MRI) and functional (e.g. sensory-evoked neural and behavioural responses, brain functional MRI) indices of development would be beneficial. We note that in the severe outcome group of dataset 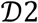, a particularly large deviation was identified at 27.3 weeks PMA (see Figure 6aii,bii). When investigating this infant’s recording further (by AD), it was confirmed that the baby was indeed very clinically unstable, with a history of seizure activity, generally suppressed baseline EEG and alternating, abnormal rhythmic activity. Further investigations into associations between the brain age delta magnitude and these contemporaneous and follow-up assessments will be highly valuable in advancing model validity and appreciating the potential clinical value of the Sinc brain age prediction model.

It is important to note that the focus of this manuscript was to provide an efficient diagnostic approach for identifying abnormal brain maturation and to additionally show that this metric correlates strongly with long term neurodevelopmental outcome. We do not, however, suggest a cause for deviations between true age and brain age (i.e. brain age deltas) in this study nor that this is directly associated to specific environmental or genetic causes. There is increasing evidence that large brain age deltas may be a symptom of pre-existing conditions from birth (such as genetic factors or low birth weight) which has a lasting impact on the infant’s development presented through alterations in brain age trajectories (Vidal-Pineiro et al., 2021). Regardless of the specific causes of brain age deltas, it is clear that the magnitudes of these deviations are of biological and clinical interest, and the ability to track and estimate brain age deviations with a model such as Sinc provide a means to identify effects as soon as they manifest potentially allowing for rapid clinical responses.

## 5. Conclusions

We outline a deep learning approach for infant brain age prediction and follow-up BSID-II outcome risk-stratification with dramatically reduced EEG data requirements relative to previous proof-of-concept studies. In an independent hold-out dataset, our Sinc model accurately predicts infant brain age and significantly distinguishes infants with normal outcome from those with severely abnormal outcome using a 1-channel bipolar montage setup and 20 min recording duration. The model also accurately predicts infant brain age when applied to data collected at an independent site. This objective and automated deep learning approach thus displays potential clinical utility for cot-side monitoring and use in neurological function assessment. A major next objective will be the efficient deployment of this model into the hospital setting using clinical grade bipolar montage data.

## Data availability statement

Due to ethical restrictions and the sensitive nature of these data, it is not possible to publicly share the supporting data.

## Code availability statement

The underlying code for the deep learning models, including the training, validation, and testing processes are openly available for download using the following GitHub link: https://github.com/amirans65/brainagemodel.

## CRediT authorship contribution statement

**Amir Ansari:** Methodology, Software, Validation, Formal analysis, Investigation, Writing – Original Draft, Visualisation. **Kirubin Pillay:** Conceptualisation, Methodology, Software, Validation, Formal analysis, Investigation, Data Curation, Writing – Original Draft, Visualisation. **Luke Baxter:** Formal analysis, Visualization, Writing – Review & Editing. **Emad Arasteh:** Formal analysis, Writing – Review & Editing. **Anneleen Dereymaeker:** Investigation, Resources, Data Curation, Writing – Review & Editing. **Gabriela Schmidt Mellado:** Visualization, Data Curation, Writing – Review & Editing. **Katrien Jansen:** Investigation, Resources, Data Curation, Writing – Review & Editing. **Gunnar Naulaers:** Resources, Writing – Review & Editing, Supervision, Funding acquisition. **Aomesh Bhatt:** Writing – Review & Editing, Supervision. **Sabine Van Huffel:** Resources, Writing – Review & Editing, Supervision, Funding acquisition. **Caroline Hartley:** Data Curation, Writing – Review & Editing, Supervision. **Maarten De Vos:** Conceptualization, Resources, Writing – Review & Editing, Supervision, Funding acquisition, Project Administration. **Rebeccah Slater:** Resources, Writing – Review & Editing, Supervision, Funding acquisition, Project Administration.

## Acknowledgements

We would like to thank all parents and infants involved in the study and staff at the UZ Leuven and John Radcliffe Hospitals who helped with data collection.

A.H.A. is supported by the FWO postdoctoral fellowship.

K.P., G.S.M, A.B., and R.S. are funded by a Senior Wellcome Research Fellowship awarded to R.S. (207457/Z/17/Z). LB is funded by a BLISS research grant.

S.V.H. and M.D.V. are funded by Bijzonder Onderzoeksfonds KU Leuven (BOF), Prevalentie van epilepsie en slaapstoornissen in de ziekte van Alzheimer [C24/18/097], Fonds voor Wetenschappelijk Onderzoek-Vlaanderen (FWO), PhD/Postdoc grants, and Agentschap Innoveren en Ondernemen (VLAIO) 150466: OSA+.

CH is funded by a Wellcome Trust/Royal Society Sir Henry Dale Fellowship (213486/Z/18/Z).

KU Leuven Stadius acknowledges the financial support of imec, EU: EU H2020 FETOPEN ‘AMPHORA’ [766456], EU H2020 MSCA-ITN-2018: ‘INtegrating Magnetic Resonance SPectroscopy and Multimodal Imaging for Research and Education in MEDicine (INSPiRE-MED)’, funded by the European Commission under Grant Agreement [813120], EU H2020 MSCA-ITN-2018: ‘INtegrating Functional Assessment measures for Neonatal Safeguard (INFANS)’, funded by the European Commission under Grant Agreement [813483], EIT 19263 – SeizeIT2: Discreet Personalized Epileptic Seizure Detection Device; Flemish Government; COST action CA20124 https://www.cost.eu/actions/CA20124/. This research also received funding from the Flemish Government (AI Research Program).

A.H.A, S.V.H. and M.D.V. are also affiliated to Leuven.AI - KU Leuven institute for AI, B-3000, Leuven, Belgium.

