## Supplementary information for "Brain age as an estimator of neurodevelopmental outcome: A deep learning approach for neonatal cot-side monitoring"

### S.1. Alternative deep learning architectures

A number of different designs were considered. Figure 2b (main text) shows a summary of these alternative architectures which include the original sequential CNN block (Figure 2bi), an extension that includes a 'residual shortcut' (Figure 2bii) (He et al., 2016), and the multi-scale Inception block with independent parallel convolutions (Figure 2biii) (Szegedy et al., 2017). In total, four additional deep learning architectures were considered and in all cases the input EEG segment structure was the same as for Sinc:

1. No FEII: Following the same design as the proposed Sinc architecture but without the entire Feature Extraction II (FEII) portion (see Figure 2a).
2. CNN: Replaces the Sinc blocks with the same CNN layer used elsewhere in the model (Figure 2bi design).
3. Residual: Similar to CNN but including the additional residual shortcut (Figure 2bii).
4. Inc: Replacing the Sinc blocks with traditional Inception blocks (Figure 2biii).

All models were trained and optimised independently and, outside of the architectural changes, all other conditions were exactly matched.

### S.2. Splitting dataset $\mathcal{D}1$ into training set and test set

Dataset $\mathcal{D}1$ was divided by recording into training and test sets of size 50 and 47 recordings, respectively. These were age-stratified by first dividing PMA into two-week intervals (27–28, 29–30, …, 42-43 weeks PMA). Each recording within an interval was then randomly assigned to either the training or test set with 50% probability, ensuring a good representation in both sets across PMA. Supplementary Figure 1 shows the PMA distribution across the training and test sets, demonstrating that the two groups were well matched. A recording-wise test-train split was chosen as it allowed a better stratification by age than splitting by infant. While a recording-wise split does not guarantee full statistical independence between the training and test sets due to recordings from a single infant possibly featuring in both sets, this issue is mitigated due to three considerations. First, the wide intervals (typically more than two weeks) between an infant's recordings limited infant-specific dependencies between recordings. Second, if there are still some dependencies in the dataset $\mathcal{D}1$, its purpose is to assess the relative improvement of training design and model architectures which is still valid.

Third, brain age prediction was subsequently assessed in datasets $\mathcal{D}2$ and $\mathcal{D}3$, directly testing and validating generalisability to independent datasets.

**S.3. Deep learning model training in dataset $\mathcal{D}1$**

To prevent over-fitting during deep learning model training, early stopping was used by assessing the change in model performance based on a validation set. The validation set was formed by removing the last 25% of each recording in the training set. This ensured that the validation set was stratified in the same way as the training set such that model updates during training were always based upon a good age representation in the data.

Two sources of Gaussian noise were added to the deep learning networks to improve robustness. First, Gaussian noise (standard deviation = 0.001) was added to the standardized input EEG. This helps the network to overcome noisy EEG and is a common approach used to prevent overfitting in deep learning models to noise in the data and has shown great success across many other deep learning applications (Audhkhasi et al., 2013; Bishop, 1995; Ghose et al., 2020; Koistinen and Holmstrom, 1991; Vincent et al., 2010; Yin et al., 2015). Injecting small random noise to the input signal helps the network learn to ignore such noisy patterns and therefore better generalise to new, unseen datasets.

Additionally, small Gaussian noise (standard deviation = 1 day) was added to the PMA target labels. As the PMAs of the recordings are sparsely scattered (and repeated for each segment in a recording), this helped the network to tolerate small prediction errors and further improve generalization performance. This was justifiable as it is known within the paediatric community that there is in fact an inherent error when determining PMAs in the clinical setting and, according to the American Academy of Paediatrics, this can vary by as much as two weeks (Engle et al., 2004). Consequently, adding Gaussian noise with a deviation of only 1 day is sufficient to generate robust training over the PMAs without affecting the correctness of the ground truth.

The EEG recording was fully segmented into contiguous 30 s segments (e.g. a 1 hr recording was segmented into 120 segments). As the durations of the recordings are not consistent, conventional segmentation into 30 s segments using a sliding window ensures that longer duration recordings are more emphasised during training resulting in a bias. To solve for this, a fixed number of segments (n=1000) is picked at random from every recording with replacement (bootstrapping) for each batch during training. This results in a total of 6.5M bootstrapped segments per training epoch.

For all deep learning models, the model produces a brain age prediction for each 30 s recording segment, and the set of estimates per recording is aggregated into a single predicted brain age value (see section 2.4.3). To partially correct for the training bias resulting from the non-uniform distribution of the data with PMA, a training weight is assigned to each segment depending on the frequency of their corresponding PMA. To calculate these training weights (or class weights) an approach used in classification tasks was employed by grouping the PMAs into ranges and calculating each weight based on the inverse harmonic mean of the number of recordings per class (King and Zeng, 2001). Consequently, segments from recordings with more common PMAs have less impact on each network update during training.

Finally, as neural network training is a non-convex problem and requires a stochastic initialisation of the parameters, each trained network is not unique. Consequently, the final performance of these trained networks varies. To achieve a robust solution, a deep ensemble approach was used by repeatedly training the model ten times using different random initialisations i.e. a 10-learner ensemble method (Fort et al., 2020).

### S.4. Interpreting Sinc model performance using dataset $\mathcal{D}1$

*Input-loss minimization*

In input-loss minimization, the input is initialised as Gaussian noise and then adapted to minimize the loss function (while keeping the trained parameters constant), for a specific target PMA. The output is thus interpreted as synthetic EEG data that the trained model has adapted (from Gaussian noise) to be characteristic of the specified target PMA. Inspired by the activation maximization visualization method (Erhan et al., 2009), stochastic gradient descent (SGD) is still used. However, the loss of the network corresponding to a target PMA is minimized instead of maximizing the output of the final activation function. Through this approach of input-loss minimization, changes to the input as optimized by the neural network may reveal potentially important physiological patterns that the network has identified to estimate the target PMA. Using this method, we generated synthetic EEG data for three postmenstrual weeks: PMA = 30, 35, and 40 weeks. These synthetic EEG outputs are qualitatively assessed based on known EEG maturational features over this age range (André et al., 2010), to facilitate interpretation of the Sinc model's functioning.

*Uniform Manifold Approximation and Projection (UMAP)*

UMAP is a nonlinear dimensionality reduction technique based on Riemannian geometry and algebraic topology (McInnes et al., 2020). To visualize high-dimensional data, UMAP is a valuable technique as it can preserve both local and global structures lending itself well to a wide variety of applications (Banville et al., 2019; Becht et al., 2018; Ordun et al., 2020; Probst and Reymond, 2020). Here, we used UMAPs to visualize the features extracted (activations) at an early (input to Feature Extraction I), middle (input to Feature Extraction II) and late (input to Regression) stage of the proposed network (see Figure 2a) to assess how different stages of the network transformed the data towards estimating brain age. This is summarised in Figure 5 and confirms how the layers are progressively improving discrimination of the ages with increased layer depth.

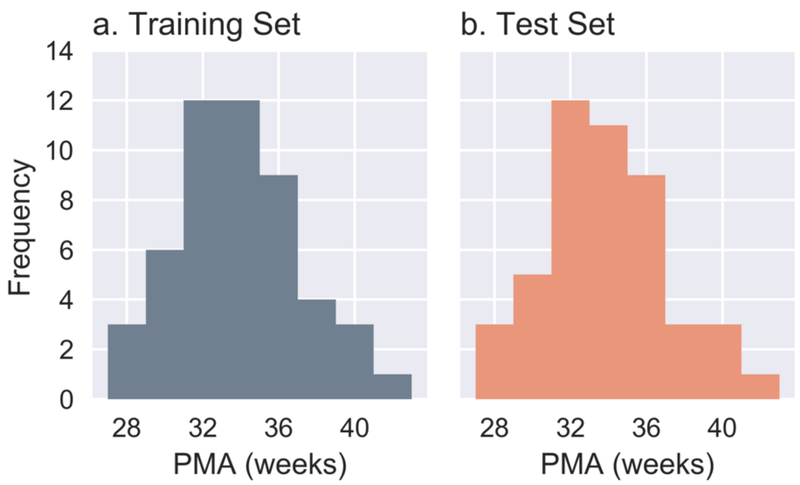


***Supplementary Figure 1. Histograms of the dataset $\mathcal{D}1$ training set and test set split of recordings by postmenstrual age (PMA).*** *Each recording was randomly assigned to either the training or test set with 50% probability, ensuring a good representation in both sets across PMA. The PMA distribution across the training and test sets thus demonstrate that the two groups were well matched.*

***Supplementary Table 1: Channel number reduction using dataset $\mathcal{D}1$.*** *Comparison of the Sinc model performance with the RF and alternative deep learning model performances. The RF model uses an 8-channel setup. The deep learning models are assessed using the same 8-channel setup and systematically reduced channel numbers (see Figure 1). Performance metrics are the MAE (in weeks), and the asterisks denote a statistically significant difference compared to the Sinc model. Values outside parentheses are for the test set (mean MAE), and values in parentheses are for the validation set (mean ± standard deviation). * = significant difference between Sinc and other models (p<0.05, paired t-test)*

| | **Number of channels** | | | |
|---|---|---|---|---|
| **Model** | **8** | **4** | **2** | **1** |
| RF | 1.01 | - | - | - |
| No FEII | 0.97* (0.71 ± 0.11) | 1.06* (0.81 ± 0.19) | 1.25* (0.91 ± 0.20) | 1.25* (0.89 ± 0.12) |
| CNN | 0.90* (0.54 ± 0.16) | 0.90* (0.57 ± 0.13) | 0.93* (0.68 ± 0.14) | 0.97* (0.67 ± 0.06) |
| Residual | 0.85* (0.53 ± 0.14) | 0.87 (0.60 ± 0.13) | 0.98* (0.71 ± 0.14) | 1.03* (0.70 ± 0.07) |
| Inc | 0.80* (0.40 ± 0.06) | 0.81 (0.55 ± 0.14) | 0.91* (0.77 ± 0.22) | 0.83 (0.70 ± 0.08) |
| **Sinc** | **0.73 (0.17 ± 0.05)** | **0.74 (0.24 ± 0.04)** | **0.77 (0.35 ± 0.05)** | **0.78 (0.50 ± 0.05)** |
